# Canine rabies detection at the livestock carcass interface: a modelling study of an integrated surveillance approach in rural India

**DOI:** 10.64898/2026.11.16.755198

**Authors:** Christine Hong, Balbir Bagicha Singh, Harsimran Singh, RS Aulakh, Garth Tarr, Jagnoor Singh Sandhu, Amrita Pattanaik, Babu N Naren, Sandul Yasobant, Ruth N. Zadoks, Victoria J. Brookes

**Affiliations:** Sydney School of Veterinary Science, Faculty of Science, The University of Sydney, Camperdown 2006, Australia; Sydney Infectious Diseases Institute, Faculty of Medicine and Health, The University of Sydney, Westmead, New South Wales, Australia; Centre for One Health, Guru Angad Dev Veterinary and Animal Sciences University, Ludhiana, Punjab 141004, India; School of Mathematics and Statistics, Faculty of Science, The University of Sydney, Camperdown, New South Wales 2006, Australia; Central Animal Research Facility, Kasturba Medical College, Manipal Academy of Higher Education, Manipal, India; Manipal Institute of Virology, Manipal Academy of Higher Education, Manipal, Karnataka 576104, India; Centre for One Health Education, Research & Development (COHERD), Indian Institute of Public Health Gandhinagar (IIPHG), Gandhinagar 382042, Gujarat, India; Section Global Health, Institute for Hygiene and Public Health (IHPH), University Hospital Bonn, Bonn 53127, Germany

**Author notes:** Corresponding author: Christine Hong. Author email addresses: Christine Hong —; Balbir Bagicha Singh —; Harsimran Singh —; RS Aulakh —; Garth Tarr —; Jagnoor Singh Sandhu —; Amrita Pattanaik —; Naren Babu N —; Sandul Yasobant —; Ruth N. Zadoks —; Victoria J. Brookes —.

**Keywords:** Active surveillance, free-roaming dogs, livestock carcass, Rabies, scenario-tree modelling

## Abstract

**Introduction:** Rabies is a deadly disease that remains endemic in many countries, particularly in rural areas. In parts of India, livestock carcass disposal sites are used by villages and visited by free-roaming dogs, providing a potential opportunity for active rabies surveillance.

**Methods:** We evaluated the potential for rabies detection at carcass disposal sites using a stochastic scenario-tree modelling framework. The model included two detection pathways: a dog scavenging pathway, in which an infectious free-roaming dog scavenges a bovine carcass and deposits detectable virus; and a bovine mortality pathway, in which the carcass originated from a rabid bovine. Model parameters were based on published literature and carcass disposal site observations. Surveillance sensitivity was estimated at carcass, district, regional, and state levels using Monte Carlo simulation assuming a 7-day surveillance period.

**Results:** Given an estimated annual canine rabies incidence risk of approximately 1,820 cases/100,000 dogs, detection probability was low at the carcass level but increased substantially across larger spatial scales. Median weekly detection probability reached 0.323 at the district level, 0.784 at the regional level, and 0.995 at the state level. A median sample size of approximately 2,220 carcasses was required to achieve 95% confidence of freedom from disease. Sensitivity analysis identified preclinical infectious duration and the number of dogs scavenging a carcass as the major drivers of surveillance performance.

**Conclusions:** These findings suggest that livestock carcass surveillance could complement existing rabies surveillance in endemic regions and help detect unrecognised circulation. High surveillance sensitivity at larger spatial scales would depend on sufficient sampling coverage across disposal sites and districts.

**Impacts:**

- Livestock carcass disposal sites could provide a complementary active surveillance interface for detecting dog-mediated rabies through both scavenging dogs and rabid bovines.
- Although detection probability was low for a single carcass, modelled 7-day surveillance sensitivity increased substantially when sampling was aggregated across districts and reached 0.995 at the state level.
- Surveillance performance was driven mainly by preclinical infectious duration and the number of dogs scavenging each carcass, highlighting the importance of broad sampling coverage and further field validation.

## 1. Introduction

Rabies is a fatal but preventable zoonotic disease that imposes a major global burden on public and animal health (Hampson et al., 2015). Although effective vaccines are available for both pre-exposure (PrEP) and post-exposure prophylaxis (PEP), rabies is still estimated to cause approximately 59,000 human deaths each year, with most cases occurring in Asia and Africa (Knobel et al., 2005, World Health Organization, 2024). The burden of rabies is not evenly distributed, with rural and resource-limited communities most affected because of limited access to PEP, diagnostic services and timely healthcare; children are also frequently affected because of their increased risk of dog-bite exposure (Hampson et al., 2008, World Health Organization, 2018). Most human rabies deaths are caused by dog-mediated transmission; therefore, controlling rabies in dog populations is central to preventing human deaths (Hampson et al., 2015).

The global strategic plan, ‘Zero by 30’, was developed in 2018 to end human deaths from dog-mediated rabies by 2030 (World Health Organization et al., 2018). This strategy requires coordinated human and animal health actions, including mass dog vaccination, improved access to PEP, public awareness, and strengthened rabies surveillance. India is central to achieving the ‘Zero by 30’ target because it has one of the highest rabies burdens globally (Sudarshan et al., 2007). India has developed a National Action Plan for Dog-Mediated Rabies Elimination by 2030, while existing national programmes also include clinical surveillance of rabies in livestock and dogs and post-mortem testing of brain tissue from animals suspected of rabies (National Centre for Disease Control, 2021, Dasgupta and Roy, 2023). However, elimination remains challenging because of the large free-roaming dog population (owned and unowned dogs that are not under direct human supervision), uneven implementation of dog vaccination and dog population management, and limited animal rabies surveillance (Pattnaik et al., 2026).

Effective rabies elimination requires surveillance systems that can identify where the virus is circulating and guide targeted control (Townsend et al., 2013). However, canine rabies surveillance in an endemic setting relies largely on the passive detection of suspect dogs, followed by sample submission and laboratory confirmation (Mananggit et al., 2021). This approach provides important information on reported cases, but it can miss local virus circulation when suspect animals are not observed, reported, sampled or tested (Minhaj et al., 2023). These limitations are likely to be greater in rural areas where veterinary services and diagnostic laboratories are less accessible (Naïssengar et al., 2021). Additional active surveillance approaches would be useful, especially if targeted to interfaces where infected dogs are likely to interact with other animals or the environment (Gill et al., 2019).

Carcass disposal sites in India could represent one such interface. In many rural villages, bovine carcasses are taken to open disposal sites (one route of carcass disposal), where they can be accessed by free-roaming dogs, other scavenging animals and insects (Singh et al., 2013). Rabies virus is transmitted in saliva, and carcasses can be sampled directly for biological material from the animal that died, including bovine saliva, while scavenging dogs might also leave canine saliva that can be sampled from the carcass (Díaz et al., 2024). Bovines (cattle and buffalo) can become infected following bites from rabid dogs, leading to mortality and disposal of carcasses (Wright et al., 2021, Vos et al., 2014). In addition, rabies cases in dogs and buffalo were found to be temporally associated in Punjab, suggesting that buffalo rabies provides an indirect signal of local dog-mediated rabies activity (Brookes et al., 2018). Because carcasses are concentrated at known disposal sites, these locations could also be repeatedly accessed for sampling. However, no published validation data were identified for the recovery of rabies virus RNA from carcass surfaces contaminated during scavenging.

Although carcass-based surveillance could provide a complementary surveillance approach, its value for detecting rabies virus at open bovine carcass disposal sites has not been assessed. It is unclear whether sampling at this interface could provide useful surveillance evidence when applied across multiple carcasses, disposal sites and districts, or whether it could be part of a freedom from disease framework, where detection probability is used to assess surveillance confidence when no positive samples are detected (Cameron, 2012, Martin et al., 2007).

The objective of this study was to determine whether bovine carcasses at carcass disposal sites could be used as surveillance sampling points for detecting rabies virus through two pathways: infectious scavenging dogs and bovines that died from rabies. Stochastic scenario-tree models were developed for both pathways and incorporated into a multi-level framework to estimate detection probabilities at the individual-animal, carcass, district, regional and state levels. A freedom-from-disease framework was used to estimate the number of carcasses required to achieve a specified level of surveillance confidence, and sensitivity analysis was used to identify the assumptions that most influenced detection probability.

## 2. Methods

The study was based on data from the state of Punjab, an agrarian state in the northwest of India. Punjab comprises 23 districts and has a total bovine population of 6,687,476, comprising farmed buffalo, farmed cattle, and stray cattle (free-roaming cattle) (Department of Animal Husbandry and Dairying, 2025, Department of Animal Husbandry and Dairying, 2023).

### 2.1. Scavenging at bovine carcass disposal sites

Data on the scavenging activity of free-roaming dogs at bovine carcass disposal sites were taken from a previous study that investigated the bovine carcass disposal system in Punjab (Singh, 2019). With one bovine carcass disposal site per one or two villages, Punjab was estimated to have 8,387 sites (Singh, 2019). In that study, seven bovine carcass disposal sites were selected, representing seven districts. Each site was monitored using continuous closed-circuit television (CCTV) surveillance for five consecutive days during four periods from July 2018 to March 2019 (20 observation days/site; total 140 site-days). Bovine carcasses that were disposed of during each 5-day period were counted. During 140 site-days, a total of 85 bovine carcasses were disposed of, giving an estimated 348,493 bovine carcasses disposed of through carcass disposal sites annually in Punjab, with a mean of 291 carcasses/district/week (Singh, 2019). The mean number of scavenging dogs recorded/carcass was 8.16 (SD 4.06; range 3–17).

### 2.2. Detection framework

The rabies detection framework considered two detection pathways: the dog scavenging pathway and the bovine mortality pathway. A stochastic scenario tree was constructed for each pathway. With respect to the dog scavenging pathway, the scenario tree described the probability that a single dog would leave detectable rabies virus in saliva on a scavenged area on a carcass. For the bovine mortality pathway, the scenario tree described the probability that the carcass originated from a bovine that had died from rabies and that detectable rabies virus would be present in its saliva. The scenario trees for these pathways are described in further detail in Section 2.2.1.

To reflect how carcass surveillance could be conducted in practice, a multi-level model was constructed with five levels: individual animal pathways, carcass-level detection, district-level detection, regional-level detection and state-level detection. The model was defined over a 7-day surveillance period, with carcass availability and sampling occurring within this timeframe. Following ascertainment of the probability of detecting rabies from the saliva left on an area scavenged by an individual dog, or from the saliva of the bovine (the individual animal pathways), the combined carcass level represented the probability of detecting rabies from one sampled carcass after combining the dog scavenging and bovine mortality pathways according to the number of scavenging dogs/carcasses. District, regional, and state levels then scaled this probability to one district, a group of 3–5 districts, and all 23 districts in Punjab, respectively.

#### 2.2.1. Scenario trees

Scenario trees were constructed using the methodology of Martin, Cameron, and Greiner (2007). Nodes in the trees represented biological processes influencing rabies detection (Figures 1 and 2). Probability distributions assigned to each node were derived from published literature or reasoned assumptions (Table 1).

**Figure 1.**
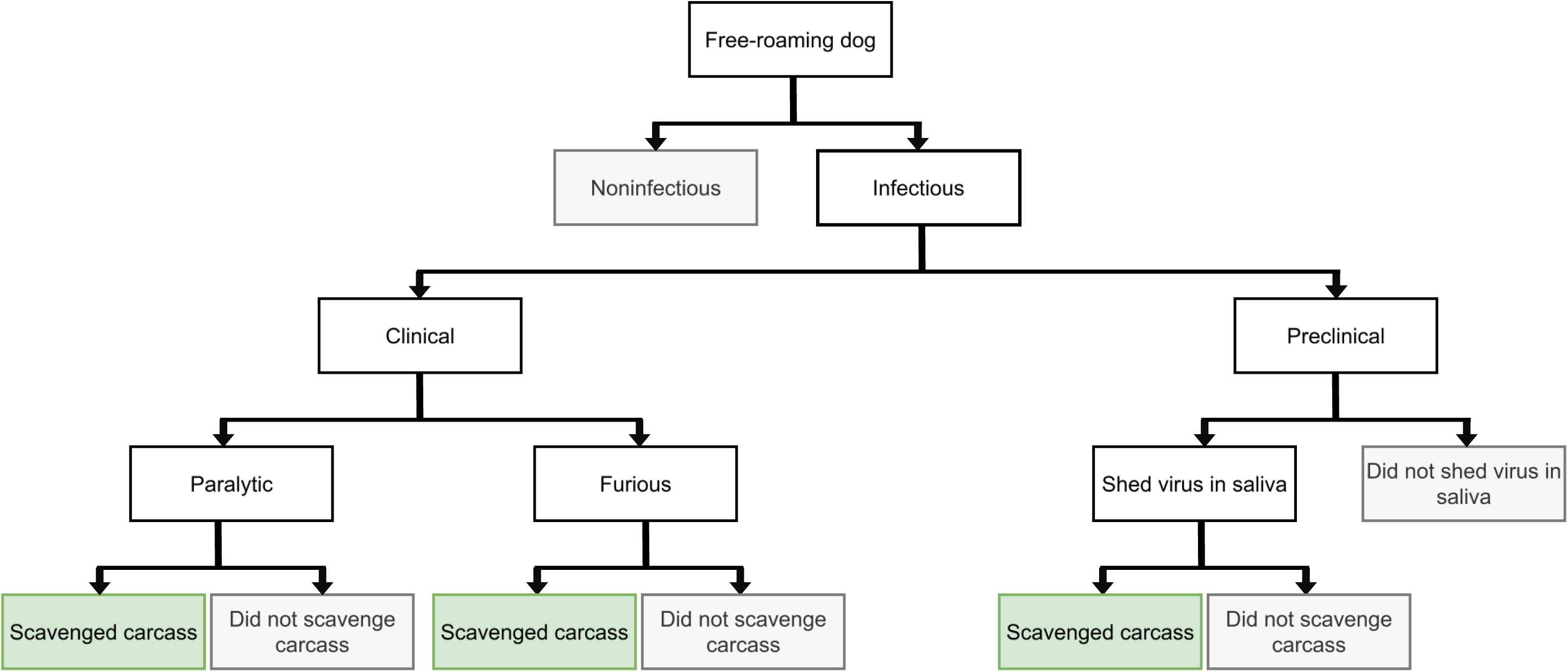
Scenario tree for the dog scavenging pathway, showing the probability of a free-roaming dog leaving rabies virus on a bovine carcass through scavenging. Green terminal nodes indicate pathways resulting in carcass scavenging and potential rabies virus detection in saliva left on the carcass.

**Figure 2.**
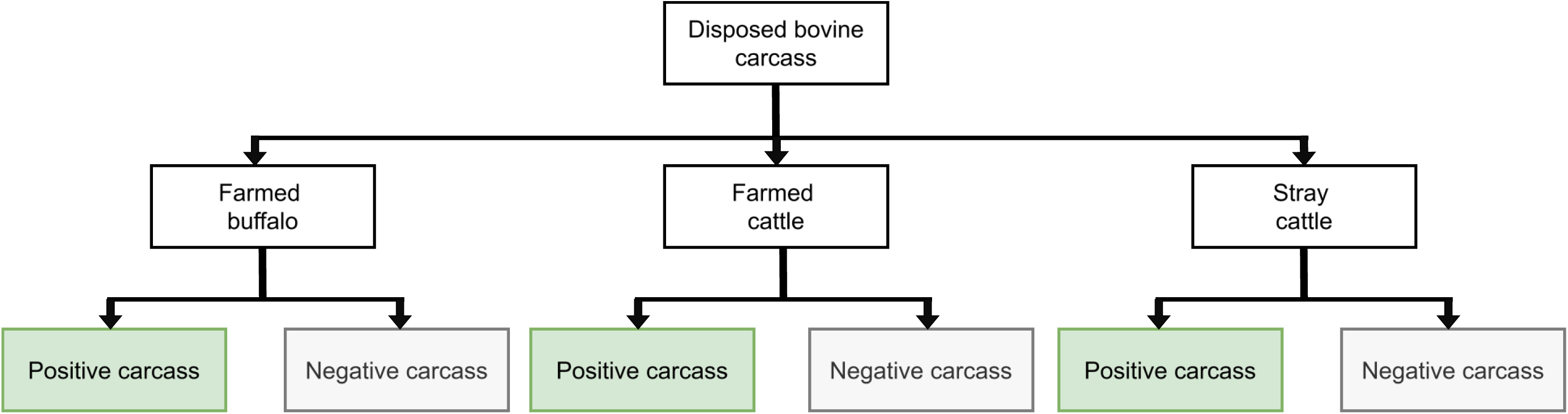
Scenario tree for the bovine mortality pathway, showing the probability that a disposed bovine carcass originated from rabies mortality. Green terminal nodes indicate rabies-positive carcasses and potential rabies virus detection in saliva from the carcass itself.

**Table 1.**
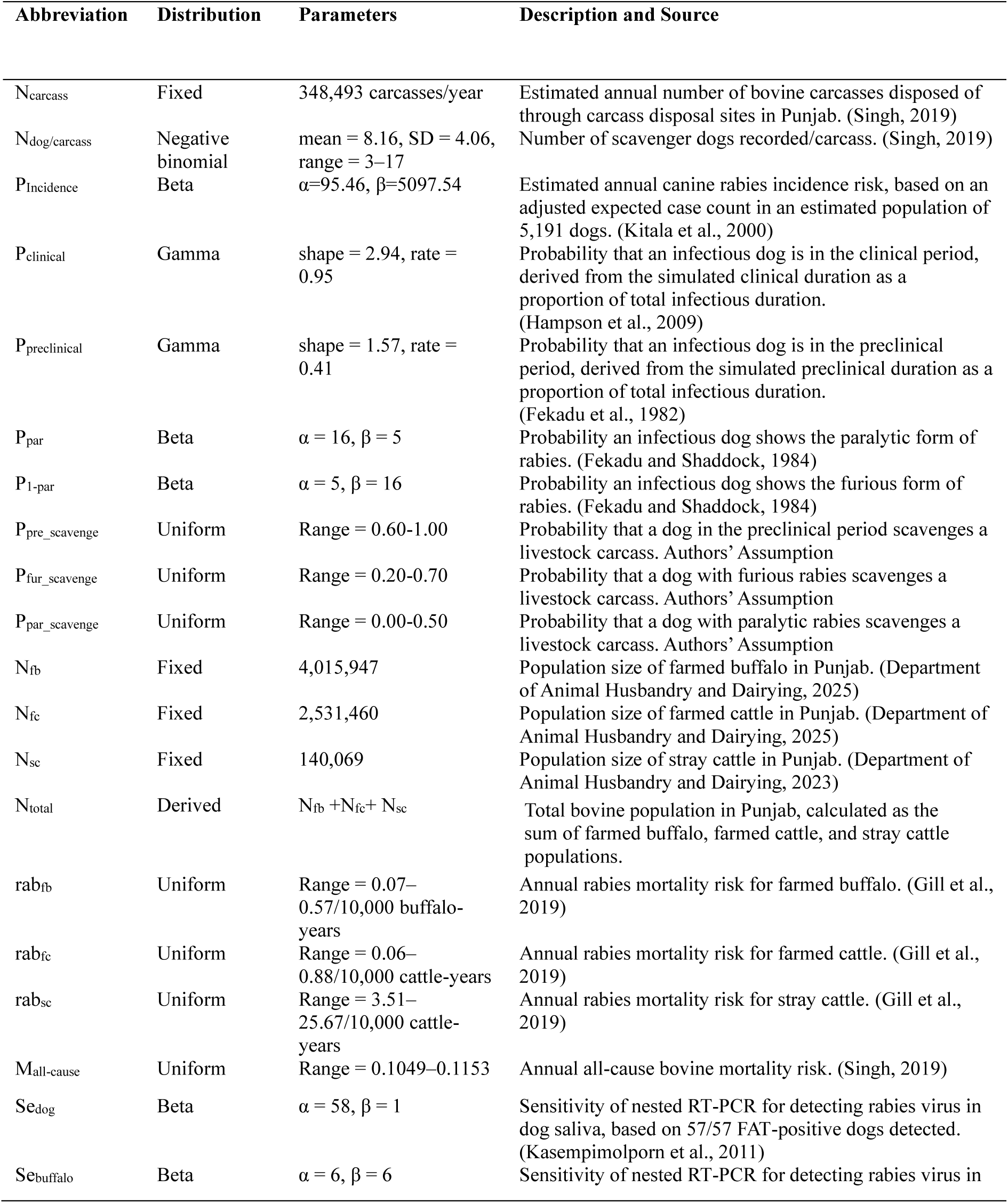
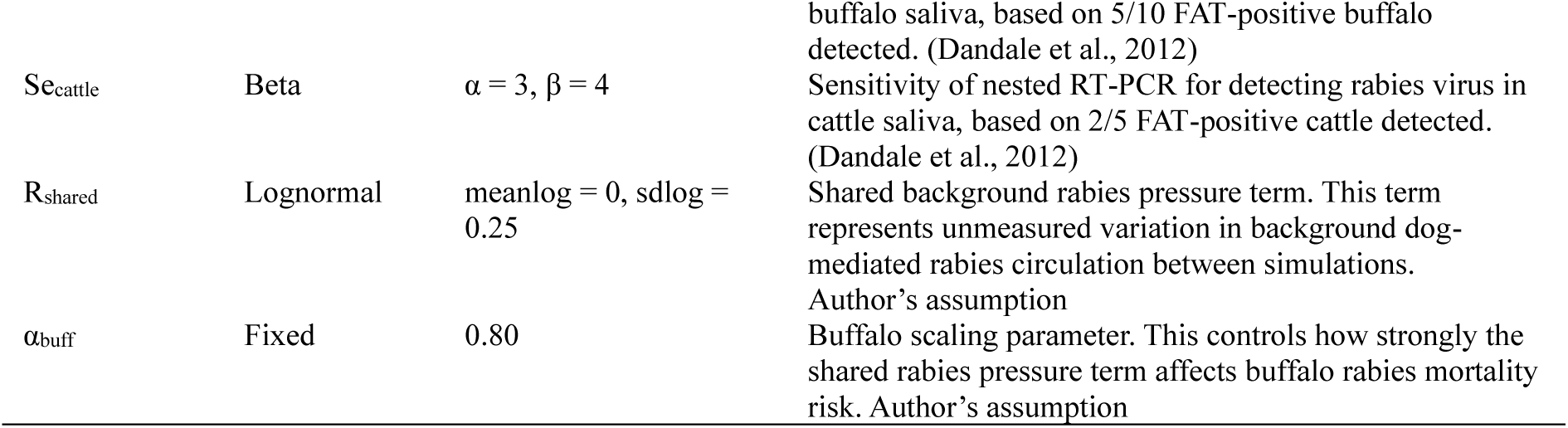
Input parameters and distributions used in the stochastic scenario tree models for the dog scavenging and bovine mortality rabies detection pathways.

In the dog scavenging pathway, the first node represented whether a free-roaming dog was rabies infectious or non-infectious (susceptible or latently infected [‘exposed’ in modelling terms]). Infectious dogs were divided into preclinical and clinical stages. During the preclinical infectious stage, dogs shed rabies virus in saliva before observable clinical signs developed. Then in the clinical stage, each dog was considered to show either the paralytic or furious form of rabies. The final node represented whether the dog scavenged the carcass, which would allow detection of rabies virus from dog saliva left on the carcass.

In the bovine mortality pathway, the first node represented the type of disposed bovine carcass, including farmed buffalo, farmed cattle and stray cattle. These groups were separated because their population size and rabies mortality risk differ. Carcasses were then classified as rabies-positive or rabies-negative according to whether they originated from rabies mortality, after which the model accounted for whether rabies virus was detectable in bovine saliva.

### 2.3. Parameter estimation

#### 2.3.1. Dog scavenging pathway

Punjab-specific canine rabies incidence estimates were available from enhanced surveillance in selected subdistricts (Gill et al., 2019). However, because these estimates were based on reported rabies-suspect animals submitted for laboratory confirmation, they were considered likely to underestimate incidence among free-roaming dogs. We could not identify studies from India that estimated overall incidence as opposed to laboratory confirmed incidence. Instead, data from an endemic setting in Kenya were used to derive an estimated annual canine rabies incidence risk of 1,820/100,000 dogs (Kitala et al., 2000). The study setting was considered relevant because dogs commonly roamed and foraged outside households, similar to free-roaming dog populations in rural India. In this Kenyan study, the total dog population across six study sublocations was estimated as 5,191 from household survey data. Rabies incidence was estimated through rabies surveillance. During the one-year surveillance period, 245 suspected dogs were captured. Of 87 dogs tested, 44 (50.6%) were confirmed to have rabies. This confirmed proportion was applied to unresolved rabies-suspect dogs (n ≈ 100), giving an adjusted estimate of 94.5 rabies cases/5,191 dogs. Annual incidence risk was then converted to a constant daily incidence rate, and daily infectious prevalence was derived from this rate and the infectious duration.

Durations were drawn from truncated gamma distributions to reflect variation across individual dogs (Fekadu et al., 1982, Hampson et al., 2009). The preclinical infectious stage had a median duration of 3.02 days (95% range: 0.30–10.77 days), and the clinical stage had a median duration of 2.75 days (95% range: 0.62–7.50 days). Both distributions were truncated at 14 days. Preclinical and clinical durations were summed to estimate the total infectious period for an individual dog in each simulation. The probability that an infectious dog was in the preclinical or clinical stage was calculated by dividing the corresponding stage duration by the total infectious period. During the clinical stage, dogs were assumed to develop the paralytic or furious form of rabies. Experimental infection outcomes indicated that, among 19 clinically rabid dogs, 15 developed paralytic rabies and 4 developed furious rabies (Fekadu and Shaddock, 1984).

The probability that an infectious dog would scavenge a carcass was parameterised using reasoned assumptions informed by literature describing scavenging and food preferences in free-ranging dogs, and the clinical presentation and behavioural changes associated with canine rabies (Bhadra et al., 2016, Sarkar et al., 2019). Preclinical dogs were assigned the highest probability of scavenging (0.60–1.00), because they were assumed to be behaving normally. Furious dogs were assigned an intermediate probability (0.20–0.70), because whilst they most likely would remain mobile, abnormal behaviour could reduce consistent feeding or scavenging. Paralytic dogs were assigned the lowest probability (0.00–0.50), because weakness, paralysis, and reduced ability to bite or swallow were expected to limit movement and scavenging.

#### 2.3.2. Bovine mortality pathway

Species- and management-specific rabies incidence rates for farmed buffalo, farmed cattle and stray cattle were obtained from an annual study conducted in Punjab, India (Gill et al., 2019) in which incidence rate was reported as laboratory confirmed cases/animal year at risk and estimated using a Poisson regression model adjusting for population size and subdistrict variation. Lower and upper incidence bounds were converted to probabilities and sampled using uniform distributions in the model.

Bovine population data were obtained from official sources based on the 20th Livestock Census. State-level farmed buffalo and farmed cattle population data were obtained from national livestock statistics, and district-level stray cattle counts were summed across Punjab (Department of Animal Husbandry and Dairying, 2025, Department of Animal Husbandry and Dairying, 2023). Subgroup-specific rabies mortality risks were weighted by population size to calculate a combined bovine rabies mortality risk. This weighting was used to estimate a single carcass-level rabies mortality risk across bovine subgroups. Annual all-cause bovine mortality risk was taken from previous work, which reported a total bovine annual mortality risk of 11% (95% CI: 10.49–11.53%) (Singh, 2019).

#### 2.3.3. Diagnostic test accuracy

Diagnostic sensitivity and specificity for saliva testing were obtained from two studies. In dogs, nested RT-PCR detected rabies virus in all 57 rabies-positive dogs and correctly classified all 180 rabies-negative dogs, giving observed sensitivity and specificity values of 100% (Kasempimolporn et al., 2011). These values were based on dogs that were mostly suspected of rabies, but they were also assumed to apply to the preclinical stage. Nested RT-PCR detected rabies viral RNA in saliva from 5 of 10 rabies-positive buffalo and 2 of 5 rabies-positive cattle, giving sensitivities of 50.0% and 40.0%, respectively (Dandale et al., 2012). All saliva samples from rabies-negative buffalo and cattle tested negative, giving an observed specificity of 100% for both species. Uncertainty in the diagnostic sensitivity was represented using the beta distributions reported in Table 1.

#### 2.3.4. Shared transmission parameter

Evidence from a previous study suggests that buffalo rabies mortality is temporally associated with local canine rabies cases, while the same pattern was not supported for cattle (Brookes et al., 2018). Therefore, a shared background rabies pressure term was included in the combined carcass-level model to represent this co-variation of dogs and buffalo within simulations (Brookes et al., 2018). The full shared term was applied as a multiplicative factor to dog infection risk. For buffalo, the effect was partially applied using a scaling parameter, αbuff = 0.80, to represent partial dependence on dog-mediated rabies pressure. Cattle were treated as independent of this shared term.

### 2.4. Model structure and implementation

A stochastic model was implemented in R version 4.5.1 (R Core Team, 2025) in which uncertainty in input parameters was incorporated using Monte Carlo simulation. Model outputs included the individual-animal pathway probabilities and rabies virus detection probabilities at the carcass, district, regional and state levels, and the number of carcasses that would need to be sampled state-wide to achieve 95% confidence of freedom from rabies. The stochastic model was run for 100,000 Monte Carlo iterations to provide precise estimates of the main summary outputs, including the mean, median, and 95% percentile intervals for carcass-, district-, regional-, and state-level detection probabilities. Sensitivity analysis was then performed using a Sobol design with 5,000 samples/input matrix and 100 bootstrap resamples.

#### 2.4.1. Level 1: Individual animal probability

The dog scavenging pathway estimated the probability that a randomly selected free-roaming dog was infectious and scavenged a bovine carcass, thereby providing an opportunity to deposit rabies virus in saliva on the carcass, *P_DogPos_*. This probability corresponded to the sum of the scavenged carcass terminal branches in the dog scenario tree. Following the scenario tree (Figure 1), *P*_Infectious_ was the probability that a free-roaming dog was infectious with rabies virus at the time of carcass contact. Infectious stages were classified as the preclinical stage *pre*, and the clinical stages in either the paralytic form *par* or furious form *fur*. For each stage, *P*(*s*) represented the probability that an infectious dog was in stage *s*, and *P*(*Scavenge* ∣ *s*) represented the conditional probability that a dog scavenged a livestock carcass given that it was in stage *s*. The stage-specific contribution, *P_DogPos_*_,*s*_, represented the probability that a randomly selected free-roaming dog was infectious, was in stage *s* and scavenged a bovine carcass:

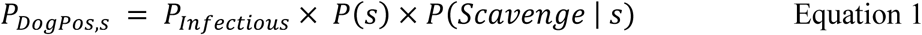

The overall probability, *P_DogPos_*, was then calculated by summing the contributions of the preclinical, paralytic and furious pathways:

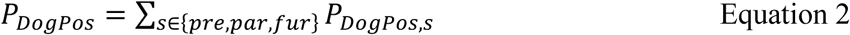

The bovine mortality pathway estimated the probability that a randomly selected disposed bovine carcass originated from a rabid bovine and therefore contained rabies virus in its saliva, *P*_BovinePos_. This was calculated according to Eq. 3 across three bovine subgroups: farmed buffalo (*fb*), farmed cattle (*fc*), and stray cattle (*sc*). For each subgroup *g*, *N_g_*represented the subgroup population size, *N*_total_represented the total bovine population, *rab_g_*represented the subgroup-specific annual rabies mortality risk, and *M*_all-cause_represented the annual all-cause bovine mortality risk. The subgroup-specific contribution, *P_BovinePos_*_,*g*_, represented the probability that a randomly selected disposed bovine carcass belonged to subgroup and originated from an animal that died from rabies:

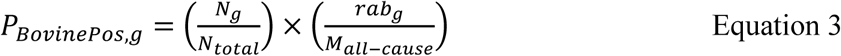

The overall bovine probability, *P_BovinePos_*, was then calculated by summing the subgroup-specific contributions:

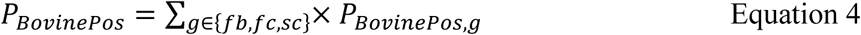

#### 2.4.2. Level 2: Carcass-level probability

This level estimated the probability that a sampled carcass was positive for rabies virus when a composite sampling protocol was applied in which for each carcass, swabs were taken from scavenged areas to detect rabies virus in dog saliva (contamination from infectious scavenging dogs) and bovine saliva (carcass originated from a rabid bovine).

The probability of detecting rabies virus from saliva deposited by infectious dogs on a sampled carcass, *P*_DogDetect_, was calculated according to Eq. 5. The number of dogs scavenging on a carcass was represented by *n_d_*. Because the source study did not distinguish single visits by multiple dogs from repeated visits by a single dog, the recorded number of scavenging dogs per carcass was treated as the number of scavenging-dog exposure opportunities in the model. The diagnostic sensitivity of PCR detection of rabies virus in dog saliva collected using a swab of scavenged areas on a bovine carcass was represented by *Se*_Dog_.

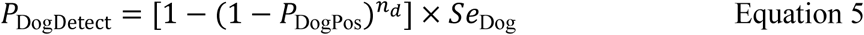

Using the subgroup-specific contributions defined in Eq. 3 and 4, the diagnostic sensitivity of PCR testing of bovine saliva was represented by *Se*_Buffalo_ for farmed buffalo and *Se*_Cattle_ for farmed and stray cattle. The probability of detecting rabies virus in bovine saliva, *P*_BovineDetect_, was calculated by applying the corresponding sensitivity to each subgroup’s contribution to *P*_BovinePos_and summing across subgroups according to Eq. 6.

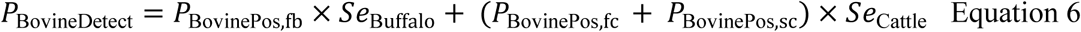

The overall probability that a carcass tested positive for rabies virus, *P*_CarcDetect_, was calculated by combining the dog scavenging and bovine mortality detection probabilities according to Eq. 7.

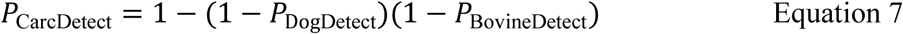

#### 2.4.3. Level 3: District-level probability

The district-level probability that at least one positive carcass was detected during a 7-day surveillance period, *P_DistrctDetect_*, was estimated according to Eq. 8. A district was defined as the administrative unit below the state level and above sub-district, village and town levels, following the administrative hierarchy used in the Census of India (Office of the Registrar General & Census Commissioner, 2022). The number of carcasses sampled in the district was represented by *n_c_*, and the carcass-level probability of detection was represented ^by^ *^P^*CarcDetect.

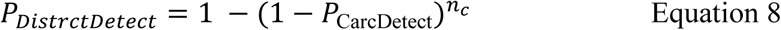

#### 2.4.4. Level 4: Regional-level probability

The regional-level probability that at least one positive carcass was detected from a group of districts sampled within the same surveillance period, *P_RegionDetect_*, was estimated according to Eq. 9. The number of districts included in the region was represented by *n*_dis_, and the district-level detection probability was represented by *P*_CarcDistrict_. Districts were assumed to be independent based on a previous analysis indicating no evidence of spatiotemporal clustering or autocorrelation in rabies cases, with a homogeneous distribution of incidence across Punjab (Brookes et al., 2018).

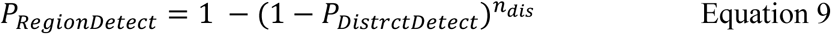

#### 2.4.5. Level 5: State-level probability

The state-level probability that at least one positive carcass was detected across all sampled districts in Punjab during a 7-day surveillance period, *P*_StateDetect_, was estimated according to Eq. 10. The total number of districts in Punjab was represented by *n*_dis_state_.

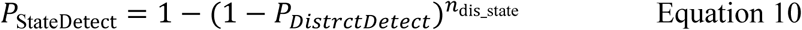

#### 2.4.6. Freedom from disease framework

In addition to estimating surveillance sensitivity, the freedom from disease framework was used to estimate how much negative carcass sampling evidence would be required to support confidence of freedom from rabies virus circulation. Eqs. 11 and 12 were used to estimate the number of sampled disposed bovine carcasses required to achieve 95% confidence of freedom from rabies virus circulation at the modelled detectable level. This was evaluated under a no-detection scenario, where no rabies-positive carcasses were detected among the sampled carcasses. The number of bovine carcasses sampled was represented by (n), and the carcass-level probability of detecting rabies virus in a sampled carcass was represented by *P*_CarcDetect_, as estimated in Level 2. To achieve 95% confidence of freedom, the probability of detecting no positives after sampling *n* carcasses was set to 0.05:

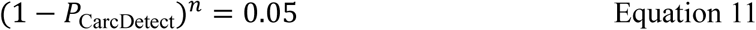

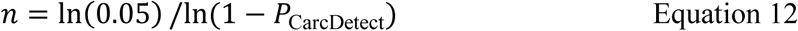

### 2.5 Sensitivity analysis

A global sensitivity analysis was conducted using the Sobol Jansen method (Jansen, 1999, Saltelli et al., 2010). Two independent Sobol design matrices were generated. Each uncertain parameter was varied across the central 95% range of its input distribution from the stochastic model (Table 1). For each parameter set, the carcass surveillance model was simulated to obtain the district-level detection probability. First-order and total-order Sobol sensitivity indices were estimated using the R package ‘sensitivity’, with bootstrap replicates used to calculate percentile intervals. First-order indices represent the proportion of output variance attributable to each parameter independently, while total-order indices represent the variance attributable to each parameter including interactions with other parameters. These indices were used to identify the parameters that contributed most to variation in the district-level detection probability.

## 3. Results

### 3.1. Individual-animal level

At an expected daily prevalence of rabies-infectious dogs of approximately 3.15 × 10⁻⁴ (equivalent to an annual canine rabies incidence risk of 1,820/100,000 dogs), the dog scavenging pathway had a total median individual-level predicted probability that a randomly selected free-roaming dog was infectious and scavenged a bovine carcass of 1.51 × 10⁻⁴ (mean: 1.87 × 10⁻⁴; 95% predicted interval [PI]: 3.02 × 10⁻⁵–5.45 × 10⁻⁴). The predicted probabilities associated with the infectious stages and clinical forms, together with the total predicted probability, are shown in Figure 3 (see Supplementary Figure S1 for the corresponding density distributions). The preclinical pathway had the highest probability, with a median of 1.20 × 10⁻⁴ (95% PI: 1.16 × 10⁻⁵–4.53 × 10⁻⁴). Lower probabilities were predicted for the furious pathway (median: 1.47 × 10⁻⁵; 95% PI: 2.51 × 10⁻⁶–5.79 × 10⁻⁵) and the paralytic pathway (median: 8.94 × 10⁻⁶; 95% PI: 3.71 × 10⁻⁷–4.06 × 10⁻⁵).

**Figure 3.**
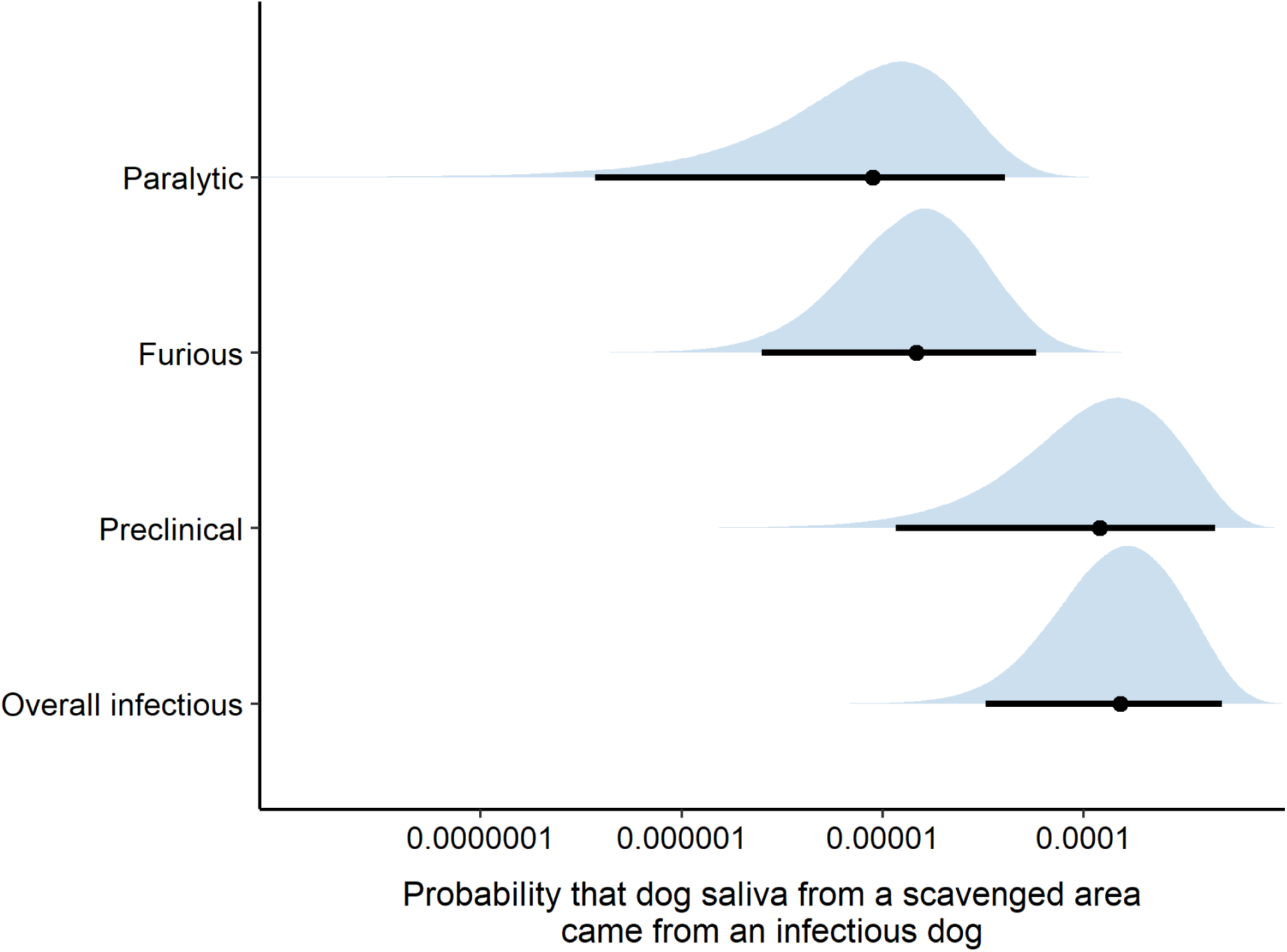
Distribution of individual-animal level probabilities that a randomly selected free-roaming dog was infectious and scavenged a bovine carcass in Punjab, by infectious stage and clinical form, including preclinical, furious, paralytic, and the combined overall pathway, given an expected daily prevalence of rabies-infectious dogs of approximately 3.15 × 10⁻⁴ (annual canine rabies incidence risk of 1,820/100,000 dogs). Dots represent median probabilities and horizontal lines represent the 95% predicted intervals.

Based on subgroup-specific annual bovine rabies mortality risks of 0.07–0.57/10,000 farmed buffalo, 0.06–0.88/10,000 farmed cattle and 3.51–25.67/10,000 stray cattle, the bovine pathway had a median individual-level probability that a randomly selected disposed bovine carcass originated from a bovine that died from rabies of 6.16 × 10⁻⁴ (mean: 6.17 × 10⁻⁴; 95% PI: 2.92 × 10⁻⁴–9.54 × 10⁻⁴). Due to the differing denominators of each scenario tree’s population (dead bovines *vs* the entire dog population), this was approximately four times higher than the median probability predicted for the dog pathway. The population-weighted subgroup predicted probabilities that a disposed carcass originated from an animal that died of rabies for farmed buffalo, farmed cattle and stray cattle, together with the overall bovine probability, are shown in Figure 4 (see Supplementary Figure S2 for the corresponding density distributions). Stray cattle had the highest median probability of 2.77 × 10⁻⁴ (95% PI: 7.70 × 10⁻⁵–4.80 × 10⁻⁴). Farmed buffalo had a median probability of 1.71 × 10⁻⁴ (95% PI: 4.31 × 10⁻⁵–3.65 × 10⁻⁴), while farmed cattle had the lowest probability among the bovine subgroups, with a median of 1.62 × 10⁻⁴ (95% PI: 2.78 × 10⁻⁵–2.97 × 10⁻⁴).

**Figure 4.**
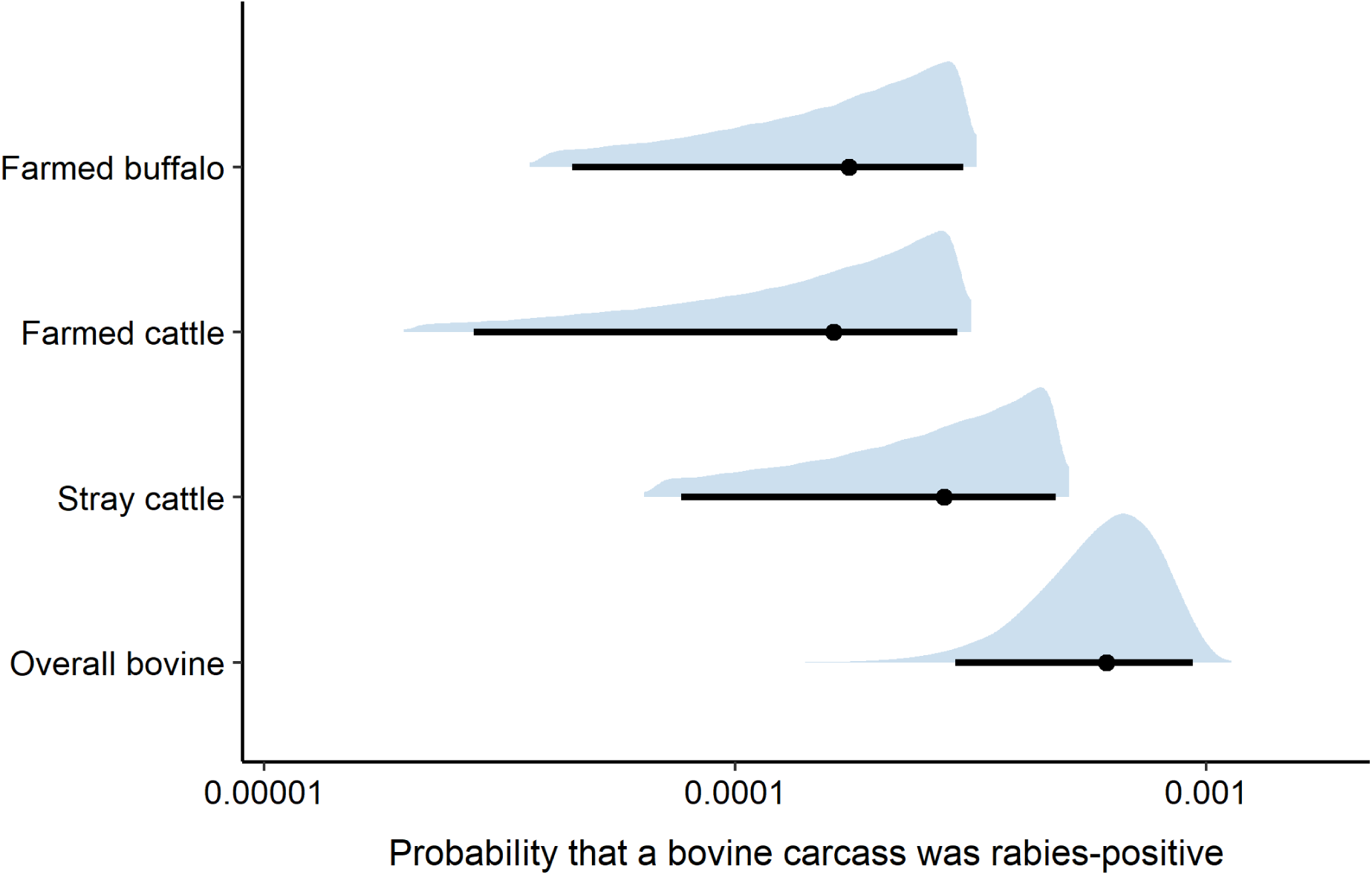
Distribution of individual-animal level predicted probabilities that a disposed bovine carcass in Punjab, India, originated from an animal that died of rabies. Farmed buffalo, farmed cattle and stray cattle estimates represent population-weighted subgroup contributions that were summed to obtain the overall bovine mortality pathway probability. Estimates were based on subgroup-specific annual bovine rabies mortality risks of 0.07–0.57/10,000 farmed buffalo, 0.06–0.88/10,000 farmed cattle, and 3.51–25.67/10,000 stray cattle. Dots represent median probabilities and horizontal lines represent the 95% predicted intervals.

### 3.2. Carcass level

The combined carcass surveillance pathway had a median carcass-level detection probability of 1.35 × 10⁻³ (mean: 1.77 × 10⁻³; 95% PI: 3.59 × 10⁻⁴–5.61 × 10⁻³). The dog scavenging pathway had a higher detection probability, with a median of 1.07 × 10⁻³ (mean: 1.50 × 10⁻³; 95% PI: 1.31 × 10⁻⁴–5.32 × 10⁻³), compared with the bovine mortality pathway, which had a median probability of 2.64 × 10⁻⁴ (mean: 2.78 × 10⁻⁴; 95% PI: 9.41 × 10⁻⁵–5.38 × 10⁻⁴). The dog scavenging pathway showed a broader distribution than the bovine mortality pathway, indicating greater variability in detection probability across simulations (Figure 5; see Supplementary Figure S3 for the corresponding density distributions). The combined carcass surveillance pathway produced the highest overall detection probability, reflecting the cumulative contributions of both surveillance pathways.

**Figure 5.**
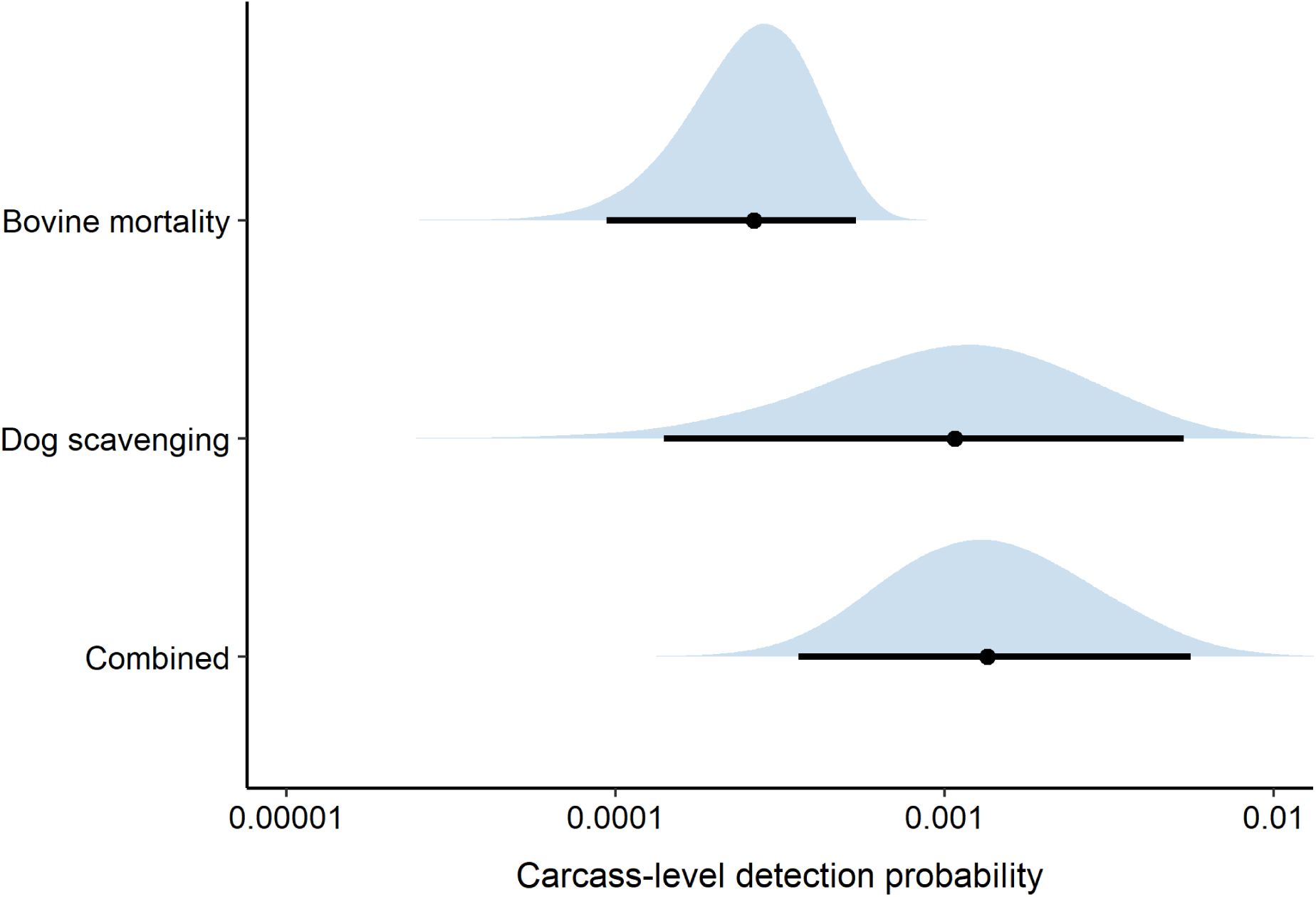
Distribution of carcass-level detection probabilities for the dog scavenging pathway, bovine mortality pathway, and the combined carcass-level surveillance pathway in Punjab, India, given an expected daily prevalence of rabies-infectious dogs of approximately 3.15 × 10⁻⁴ (annual canine rabies incidence risk of 1,820/100,000 dogs) and subgroup-specific annual bovine rabies mortality risks of 0.07–0.57/10,000 farmed buffalo, 0.06–0.88/10,000 farmed cattle, and 3.51–25.67/10,000 stray cattle. Dots represent median probabilities and horizontal lines represent 95% percentile intervals. The combined pathway represents the overall carcass-level probability of rabies detection across both surveillance pathways.

### 3.3. District, regional and state levels

Under the assumed sampling intensity of 291 carcasses/district/week, surveillance sensitivity increased from the district level to the regional and state levels (Figure 6; see Supplementary Figure S4 for the corresponding density distributions). At the district level, the probability of detecting at least one rabies-positive carcass in a 7-day surveillance period had a median of 3.24 × 10⁻¹ (mean: 3.63 × 10⁻¹; 95% PI: 9.86 × 10⁻²–8.06 × 10⁻¹). At the regional level, the detection probability increased to a median of 7.85 × 10⁻¹ (mean: 7.49 × 10⁻¹; 95% PI: 3.17 × 10⁻¹–9.99 × 10⁻¹). Under this sampling design, when surveillance was expanded to all 23 districts in Punjab, the state-level detection probability was very high, with a median approaching 1.00 (mean: 9.90 × 10⁻¹; 95% PI: 9.08 × 10⁻¹–1.00).

**Figure 6.**
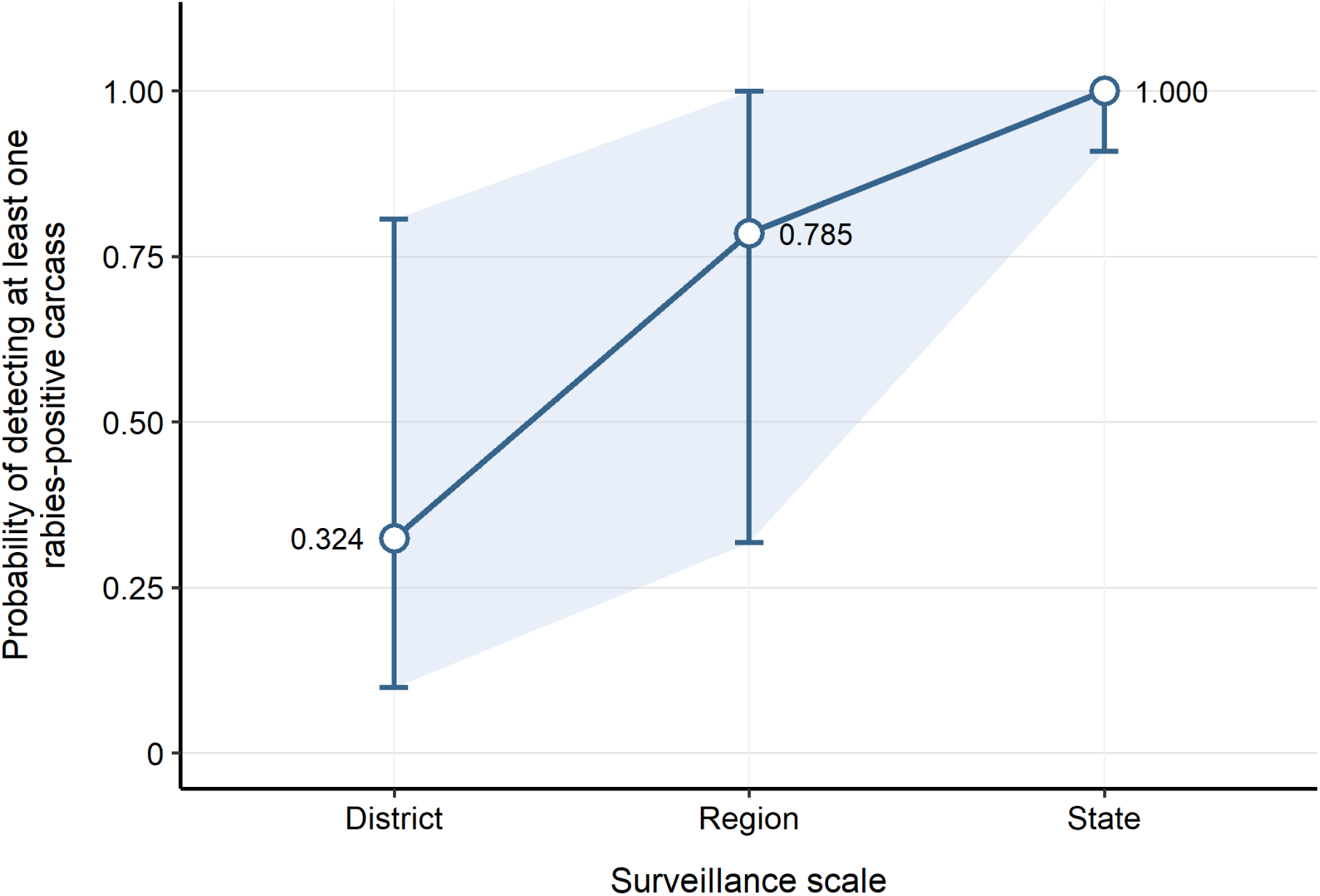
Spatial scaling of surveillance sensitivity from district to regional and state levels. Points represent median probabilities of detecting at least one rabies-positive carcass in a 7-day surveillance period in Punjab, India, given an expected daily prevalence of rabies-infectious dogs of approximately 3.15 × 10⁻⁴ (annual canine rabies incidence risk of 1,820/100,000 dogs) and subgroup-specific annual bovine rabies mortality risks of 0.07–0.57/10,000 farmed buffalo, 0.06–0.88/10,000 farmed cattle, and 3.51–25.67/10,000 stray cattle. Vertical lines represent the 95% percentile intervals derived from Monte Carlo simulations. District-level estimates assumed an average of 291 carcasses/district/week; regional- and state-level estimates were derived by scaling this across 3–5 districts and all 23 districts in Punjab, respectively.

### 3.4. Freedom from disease requirements

Figure 7 shows the distribution of the number of carcasses required to demonstrate freedom from rabies virus circulation at 95% confidence under the combined carcass surveillance framework and the modelled annual rabies risk level. Under a no-detection scenario, the estimated median number of carcasses required was 2,220 carcasses (mean: 2,793 carcasses; 95% PI: 533–8,333 carcasses; see Supplementary Figure S5 for the sample-size distribution).

**Figure 7.**
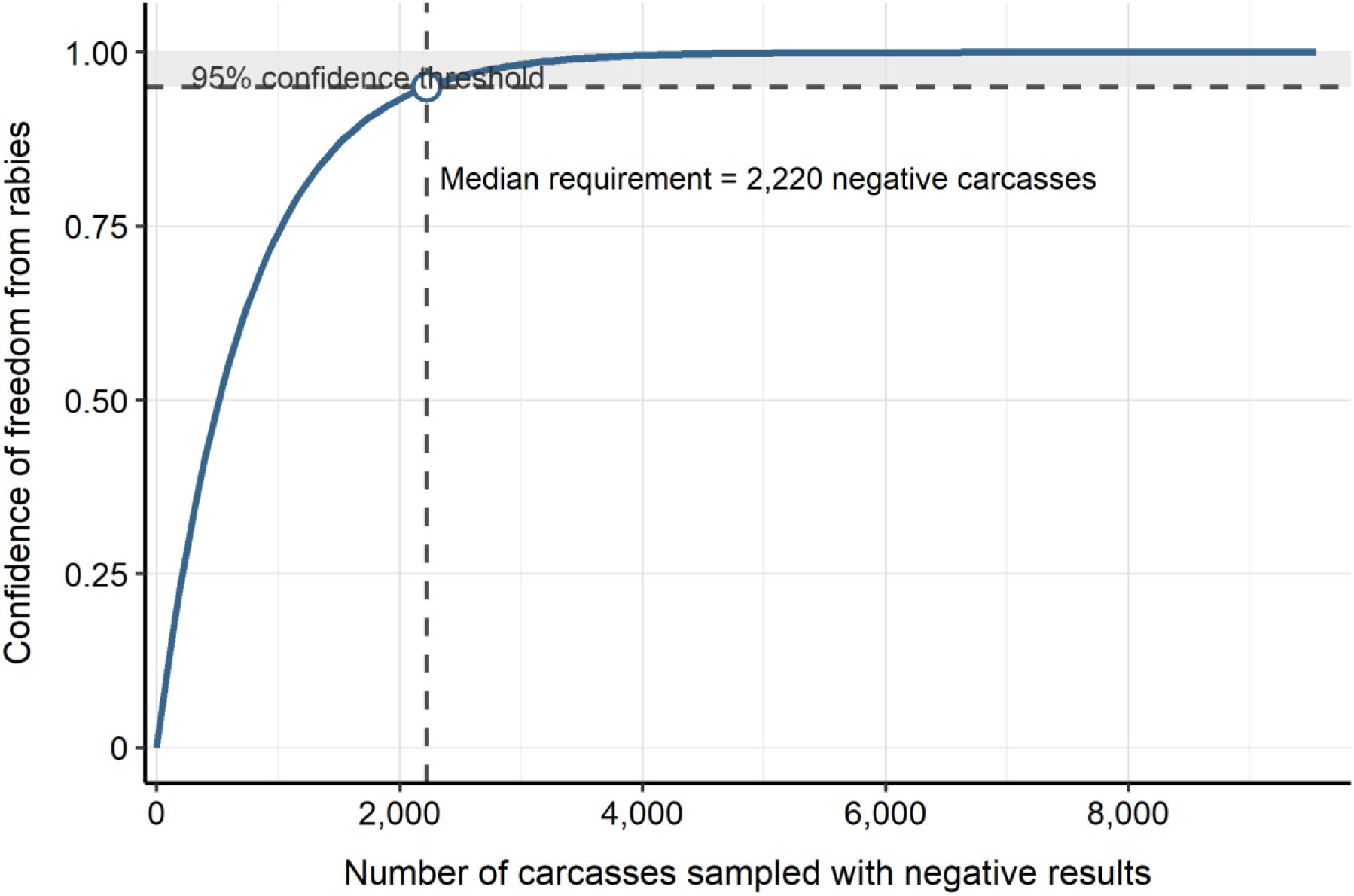
Confidence of freedom from rabies based on negative carcass surveillance under the combined carcass surveillance framework, given an expected daily prevalence of rabies-infectious dogs of approximately 3.15 × 10⁻⁴ (annual canine rabies incidence risk of 1,820/100,000 dogs) and subgroup-specific annual bovine rabies mortality risks of 0.07–0.57/10,000 farmed buffalo, 0.06–0.88/10,000 farmed cattle, and 3.51–25.67/10,000 stray cattle. The curve shows the increase in confidence of freedom as the number of carcasses sampled with negative test results increases. The dashed horizontal line represents the 95% confidence threshold, and the dashed vertical line indicates the median number of negative carcasses required to achieve this threshold.

### 3.5. Sensitivity analysis results

Figure 8 shows the first-order and total-order Sobol sensitivity indices for district-level detection probability under the combined carcass surveillance framework, with the full results provided in Supplementary Table S1. Variation in model output was mainly driven by a small number of parameters. The duration of the preclinical infectious stage had the greatest influence, with a first-order Sobol index of 0.475 and a total-order index of 0.513. This was followed by the number of dogs scavenging each carcass, with first-order and total-order indices of 0.325 and 0.363, respectively. Shared rabies pressure also contributed to model variance, with a first-order index of 0.084 and a total-order index of 0.095, although its influence was smaller than those of the two dominant dog-mediated parameters.

**Figure 8.**
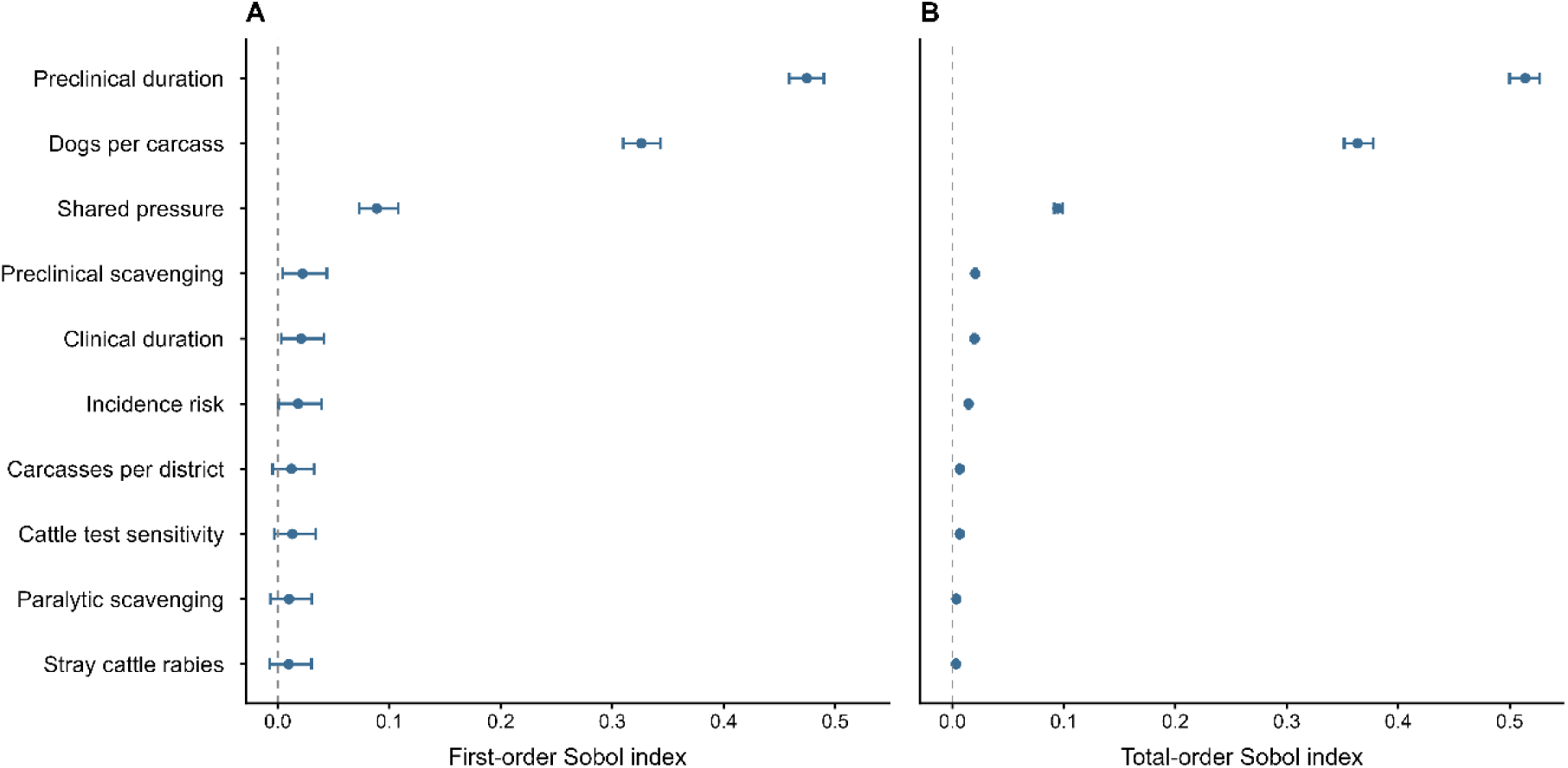
Global sensitivity analysis of the district-level surveillance model using the top 10 parameters ranked by total-order Sobol sensitivity index. Panel A shows first-order Sobol indices and Panel B shows total-order Sobol indices. Points represent Sobol index estimates and horizontal bars represent 95% bootstrap confidence intervals.

All remaining parameters had relatively small effects, with total-order Sobol indices below 0.025. Preclinical scavenging probability and clinical infectious duration each had total-order indices of approximately 0.020. Annual canine rabies incidence risk had a total-order index of 0.014. Cattle diagnostic sensitivity and the number of carcasses available within a district each had total-order indices of approximately 0.007, while all other parameters had total-order indices below 0.004. Several first-order indices were slightly negative, reflecting numerical estimation error rather than negative contributions to model variance. Total-order indices were greater than the corresponding first-order indices for the most influential parameters, indicating interaction effects within the model. Overall, model behaviour was primarily driven by preclinical infectious duration and the number of scavenging dogs per carcass.

## 4. Discussion

This study suggests that bovine carcasses at open disposal sites could be used as a tool for canine rabies surveillance, complementing existing passive rabies surveillance. A key finding from this model was that, under an estimated annual canine rabies incidence risk of 1,820/100,000 dogs, at the state level, the probability of detecting at least one rabies-positive carcass was almost certain under the assumptions used in the model. Existing national programmes in India, such as the National Action Plan for Dog-Mediated Rabies Elimination and the Integrated Disease Surveillance Programme, include clinical surveillance of rabies in livestock and dogs, as well as post-mortem testing of brain tissue from animals suspected of rabies (National Centre for Disease Control, 2021). Our framework provides a more systematic approach that does not rely on passive reporting of possible cases or dead animals. By helping identify areas where dog-mediated rabies is circulating, livestock carcass-based surveillance could support targeted control activities (World Health Organization et al., 2018).

The freedom-from-disease analysis estimated that approximately 2,220 carcasses would need to test negative to achieve 95% confidence that rabies was not circulating at or above the specified incidence risk. Even though we estimate > 6,500 carcass disposals/week in Punjab, reaching this level of aggregated sampling in practice would require coordinated surveillance across districts and sectors. It should also be noted that this estimate was based on carcass disposal sites from only seven districts in Punjab and might not represent carcass availability across all districts (Singh, 2019). Some districts might have fewer carcasses available for sampling because of differences in bovine population size or in carcass disposal practices. Nevertheless, because livestock carcasses are concentrated at established disposal sites, these sites provide practical locations for ongoing sampling, increasing opportunities for rabies detection. Sampling across multiple districts might improve detection of spatially dispersed rabies circulation compared with sampling within a single district, particularly in rural areas where animal rabies cases are less likely to be recognised or reported through passive surveillance (Gill et al., 2019, Rysava et al., 2022). Future research on carcass disposal frequency and the number of disposal sites in different areas of Punjab could improve estimates of the number of carcasses available for surveillance.

In addition to the number of carcasses required, factors that could affect detection under field conditions were not fully represented in the model and would need to be considered. In practice, bovine saliva could be sampled soon after notification of a newly deposited carcass, whereas sampling for canine saliva would need to occur after dogs had interacted with the carcass. Multiple dogs could scavenge the same carcass, and these dogs could be reluctant to provide access to people and become aggressive. This has practical implications for field team safety, including rabies vaccination and appropriate titre monitoring, and could also delay or prevent timely sampling after dog contact, reducing opportunities for detection. The previous Punjab study did not distinguish unique dogs from repeated visits by the same dogs (Singh, 2019). If repeated visits were treated as different dogs, this could increase the modelled probability that at least one scavenging dog was infectious. However, repeated visits by an infectious dog could also increase opportunities for saliva deposition and detection. As the number of scavenging dogs per carcass was influential in the sensitivity analysis, distinguishing unique dogs from repeat visits would help reduce uncertainty in this pathway. Competition around carcasses could also result in bites between dogs, potentially making carcass disposal sites foci for rabies transmission. The extent to which transmission occurs at these sites is currently unknown. If infectious dogs are clustered around particular disposal sites, or within dog populations using the same sites, surveillance sensitivity might depend not only on the number of carcasses sampled but also on the spatial distribution of the disposal sites surveyed. Sampling across multiple sites would help provide representative coverage and improve independence between samples. Future camera-trap studies could help distinguish unique dogs from repeat visits using individual visual characteristics and repeated sightings as well as investigate dog behaviour at sites.

Bovine rabies in India largely reflects spillover from the dog-mediated rabies cycle (Nagarajan et al., 2006). Livestock that roam around villages have more opportunities to encounter free-roaming dogs, which is consistent with the relatively high rabies incidence reported in stray cattle (Gill et al., 2019). Previous work in Punjab also found an association between rabies cases in dogs and buffalo (Brookes et al., 2018). Although cattle rabies is also expected to result largely from exposure to rabid dogs, the previous Punjab analysis did not identify a statistical association between reported dog and cattle rabies cases. Cattle were therefore treated as independent of the shared rabies-pressure term in the model, reflecting the available epidemiological evidence rather than an assumption that cattle rabies is unrelated to dog-mediated transmission. This remains an important evidence gap that should be addressed in future research.

The dog rabies incidence risk used in the model was based on active surveillance data from Kenya because available data from Punjab were primarily derived from passive surveillance and were likely to underestimate true incidence (Kitala et al., 2000, Gill et al., 2019). Dogs in the Kenya study were commonly allowed to roam and forage outside the household, similar to dog populations described in rural India (Tiwari et al., 2019, Kitala et al., 2000). Whilst this could be considered a limitation of the model, the incidence estimate was used to evaluate surveillance performance at a specified level of rabies incidence rather than to represent the exact incidence of canine rabies in Punjab. The model could therefore be updated with local incidence data where available.

Another limitation to consider is that the diagnostic test sensitivity estimates were obtained from saliva collected directly from animals suspected of rabies under clinical or necropsy conditions (Dandale et al., 2012, Kasempimolporn et al., 2011), whereas carcass-based surveillance would involve swabbing carcasses at disposal sites. Field sensitivity could be lower because saliva deposition might be uneven and viral material might be affected by drying, decomposition, sunlight, high temperature, and delays before sampling (McElhinney et al., 2014, Matouch et al., 1987). The timing and feasibility of these different sampling approaches would need to be evaluated under field conditions. A lateral flow assay (LFA) could provide a rapid and practical screening option at carcass disposal sites. Reported sensitivity for saliva-based LFAs has been relatively high in post-mortem samples, at 93.0% and 89.5%, while specificity was lower at 94.4% and 93.9%, respectively (Kasempimolporn et al., 2011, Bhalke et al., 2026). Although specificity was also relatively high, lower specificity can lead to more false-positive results when screening large numbers of carcasses, particularly when rabies prevalence is low (Léchenne et al., 2016, Trevethan, 2017). LFA performance would therefore need to be evaluated on carcass samples under field conditions before large-scale surveillance. Evidence for rabies virus detection in bovine saliva was also based on small numbers of animals, adding further uncertainty to test performance (Maxim et al., 2014, Dandale et al., 2012, Delpietro et al., 2001, Dandale et al., 2013).

## 5. Conclusion

The probability of detecting rabies virus using bovine carcasses (bovine saliva, canine saliva) as the sample source increased substantially as sampling was aggregated from a single district to all 23 districts at the state level within a 7-day surveillance period. Many negative samples would be needed to provide high confidence of freedom from canine rabies virus transmission in Punjab, but carcass-based surveillance could be used as an active surveillance tool if supported by sufficient state-wide organisational capacity, reinforcing campaigns for ‘Zero by 30’ and helping to identify higher-risk locations where dog vaccination, public awareness, and other rabies control activities could be targeted (World Health Organization et al., 2018). Further field data are needed to test this approach in practice, including dog scavenging behaviour, carcass sampling methods, and diagnostic test performance under field conditions.

## Author Contributions

**Conceptualization:** C.H., V.J.B., R.N.Z.; **Methodology:** C.H., V.J.B., R.N.Z.; **Formal analysis:** C.H.; **Validation:** V.J.B., G.T.; **Supervision:** V.J.B., R.N.Z.; **Writing – original draft:** C.H.; **Writing – review and editing:** all authors.

## Ethics Statement

Ethical approval was not required because this study did not involve human participants or live animals.

## Data Availability Statement

The data used to parameterise the model are derived from published sources and are provided within the article and its Supporting Information. No original empirical data were generated for this study.

## Supporting information

Supplementary material. Click Upload Files

