## Supplementary material. Click Upload Files for "Canine rabies detection at the livestock carcass interface: a modelling study of an integrated surveillance approach in rural India"

**Supplementary Materials**

### Table S1. Sobol first-order and total-order sensitivity indices

| **Parameter** | **R model name** | **First-order index** | **Total-order index** |
| --- | --- | --- | --- |
| Preclinical infectious duration | q_preclinical_duration | 0.474560 | 0.513952 |
| Clinical infectious duration | q_clinical_duration | 0.020790 | 0.019933 |
| Annual canine rabies incidence risk | Incidence_risk | 0.018229 | 0.014505 |
| Paralytic proportion given clinical disease | P_paralytic_given_clinical | 0.008436 | 0.002693 |
| Preclinical scavenging probability | q_scavenge_pre | 0.022178 | 0.020502 |
| Furious scavenging probability | q_scavenge_fur | 0.007587 | 0.001526 |
| Paralytic scavenging probability | q_scavenge_par | 0.010023 | 0.003588 |
| Number of dogs on carcass | nd_dogs_on_carcase | 0.326063 | 0.363491 |
| Dog diagnostic sensitivity | Se_dog | 0.007202 | 0.000413 |
| Buffalo diagnostic sensitivity | Se_buffalo | 0.007096 | 0.000738 |
| Cattle diagnostic sensitivity | Se_cattle | 0.013017 | 0.006750 |
| Farmed cattle rabies risk | rab_fc | 0.007698 | 0.001434 |
| Farmed buffalo rabies risk | rab_fb | 0.007457 | 0.001823 |
| Stray cattle rabies risk | rab_sc | 0.009772 | 0.003216 |
| All-cause bovine mortality | mortality | 0.006921 | 0.000066 |
| Shared rabies pressure | q_shared_pressure | 0.088741 | 0.094505 |
| Number of carcasses per district | n_carcases_district | 0.012214 | 0.006773 |

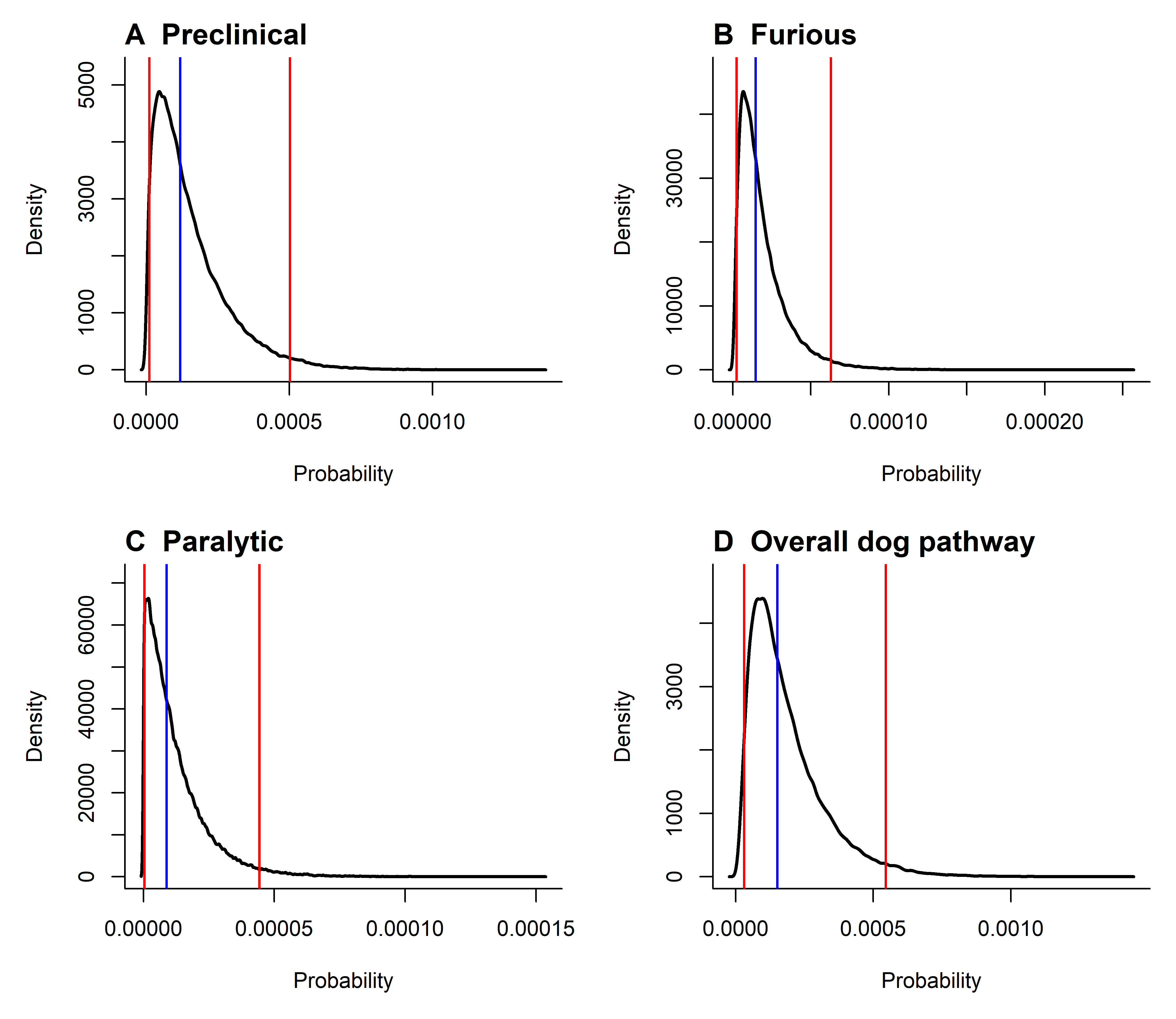

Figure S1. Density distributions of the individual-animal level predicted probabilities for the dog scavenging pathway. The preclinical, furious and paralytic panels show the probability that a randomly selected free-roaming dog was infectious, was in the corresponding infectious stage or clinical form, and scavenged a bovine carcass. The overall dog pathway represents the probability that a randomly selected free-roaming dog was infectious and scavenged a bovine carcass. The solid vertical line indicates the median and dashed vertical lines indicate the 2.5th and 97.5th percentiles.

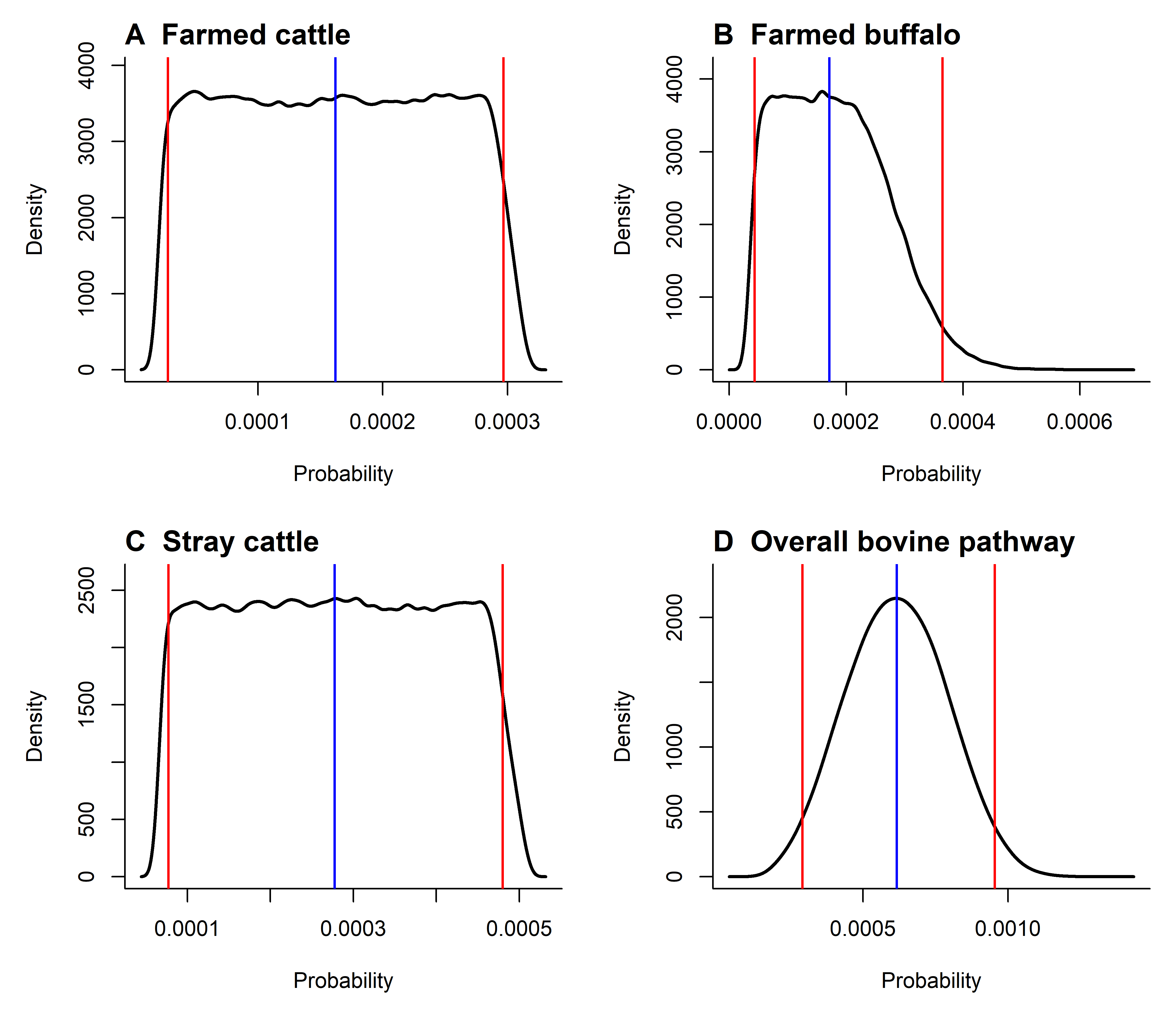

Figure S2. Density distributions of the individual-animal level predicted probabilities for the bovine mortality pathway. Farmed cattle, farmed buffalo and stray cattle represent the population-weighted probabilities that a randomly selected disposed bovine carcass belonged to the corresponding subgroup and originated from an animal that died from rabies. The overall bovine pathway represents the probability that a randomly selected disposed bovine carcass originated from an animal that died from rabies. The solid vertical line indicates the median and dashed vertical lines indicate the 2.5th and 97.5th percentiles.

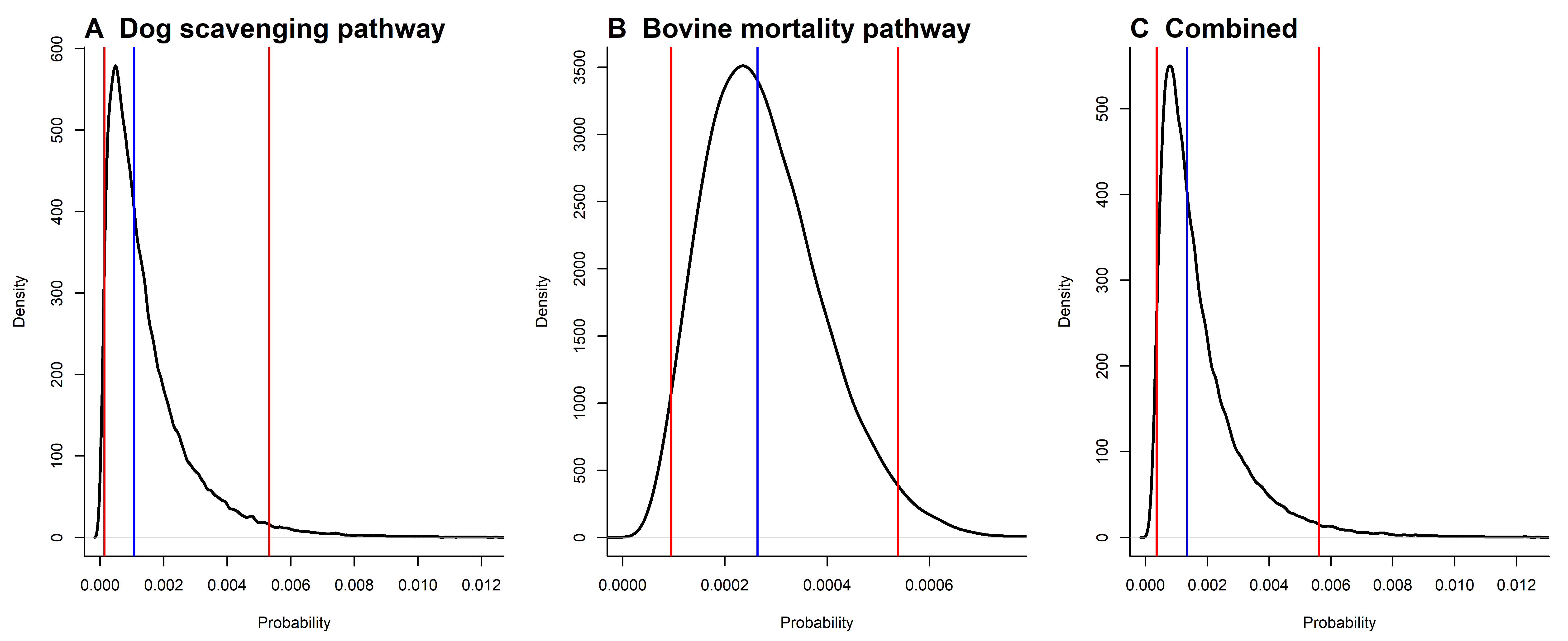

Figure S3. Density distributions of the carcass-level probabilities of detecting rabies virus from one sampled carcass through the dog scavenging pathway, bovine mortality pathway and the combined carcass surveillance pathway. The combined pathway represents the overall probability of detecting rabies virus from one sampled carcass through either pathway. The solid vertical line indicates the median and dashed vertical lines indicate the 2.5th and 97.5th percentiles.

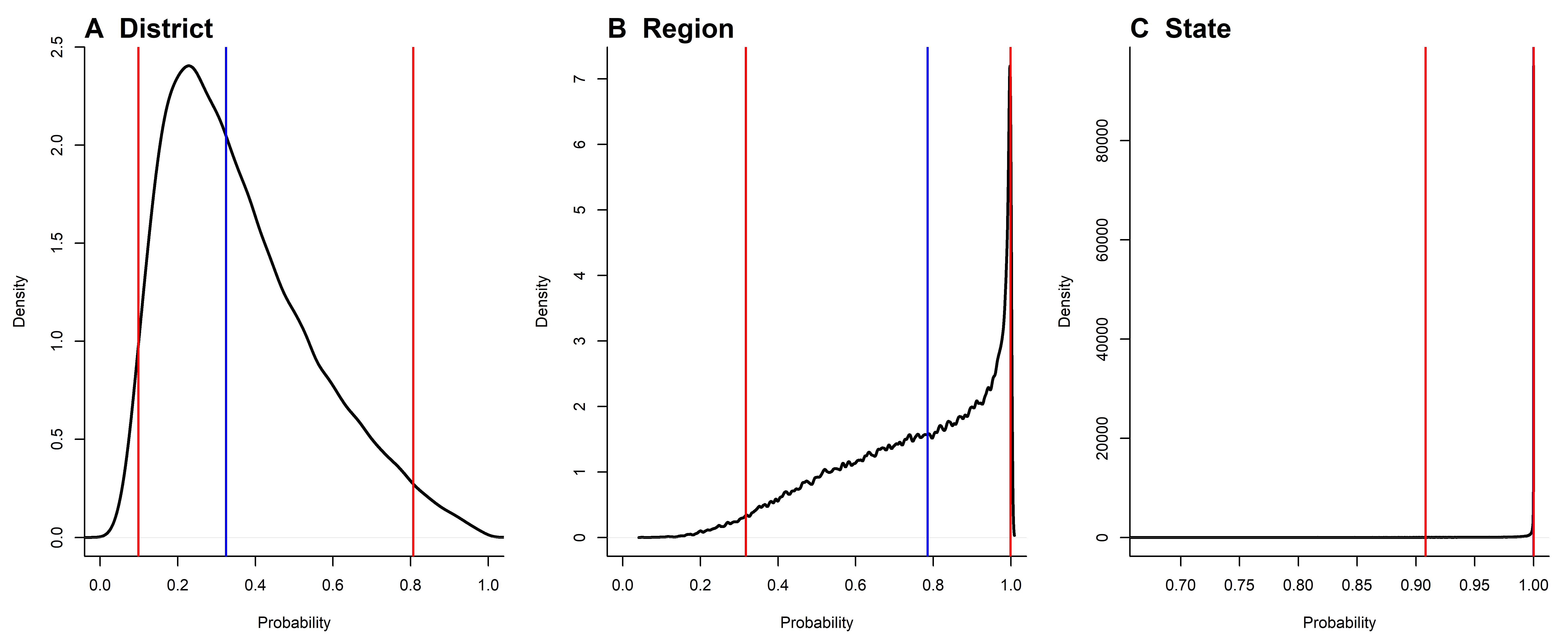

Figure S4. Density distributions of the probability of detecting at least one rabies-positive carcass during a 7-day surveillance period at district, regional and state levels. District-level estimates assumed sampling of 291 carcasses/district/week, with regional and state estimates scaled across 3–5 districts and all 23 districts in Punjab, respectively. The solid vertical line indicates the median and dashed vertical lines indicate the 2.5th and 97.5th percentiles.

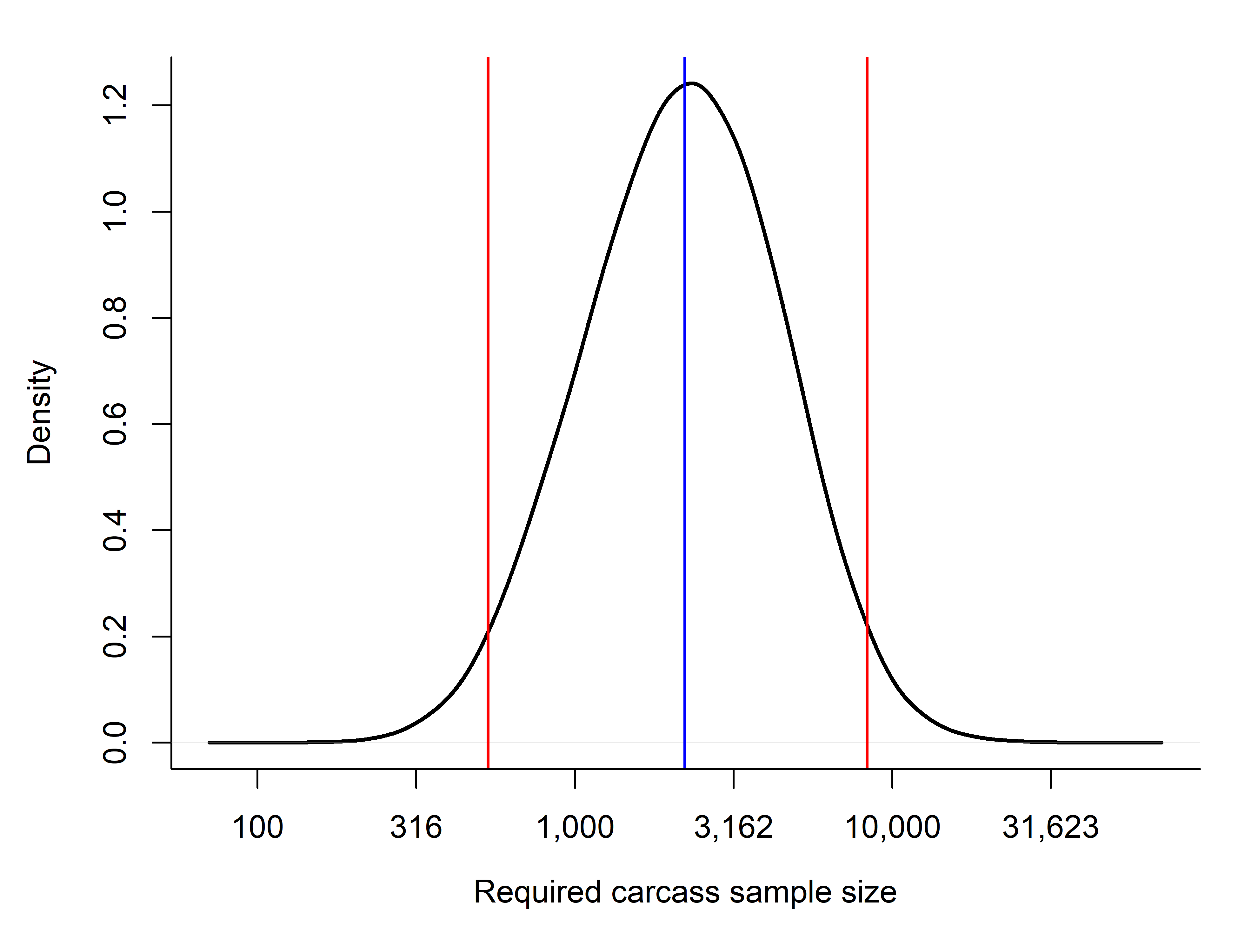

Figure S5. Distribution of the number of negative carcass samples required to achieve 95% confidence of freedom from rabies virus circulation at the specified detectable level. A log-scaled horizontal axis is used to display the right-skewed distribution. The solid vertical line indicates the median and dashed vertical lines indicate the 2.5th and 97.5th percentiles.
